# Mutations Leading to Ceftolozane/Tazobactam and Imipenem/Cilastatin/Relebactam Resistance During *in vivo* exposure to Ceftazidime/Avibactam in *Pseudomonas aeruginosa*

**DOI:** 10.1101/2024.09.10.612322

**Authors:** Glenn J. Rapsinski, Alecia B. Rokes, Daria Van Tyne, Vaughn S. Cooper

## Abstract

Identifying resistance mechanisms to novel antimicrobials informs treatment and antimicrobial development, but frequently identifies multiple candidate resistance mutations without resolving the driver mutation. Using whole genome sequencing of longitudinal *Pseudomonas aeruginosa* that developed imipenem/cilastatin/relebactam and ceftolozane/tazobactam resistance during ceftazidime/avibactam treatment, we determined mutations resulting in cross-resistance. Penicillin-binding protein *ftsI*, transcriptional repressor *bepR*, and virulence regulator *pvdS* were found in resistant isolates. We conclude that peptidoglycan synthesis gene mutations can alter the efficacy of multiple antimicrobials.

---

Antimicrobial resistance (AMR) is a public health crisis costing over a million lives and billions of dollars in healthcare costs each year (1, 2). Cross resistance, resistance to multiple antimicrobials from a single bacterial adaptation, further complicates AMR (3). Cross resistance can occur through non-specific mechanisms such as multi-drug efflux (4), and by class-specific mechanisms (5, 6). Nosocomial pathogens, like *Pseudomonas aeruginosa*, are primarily responsible for the AMR burden (2). *P. aeruginosa*, a Gram-negative opportunistic pathogen, can cause ventilator-associated pneumonia, burn infections, sepsis, and chronic infection in people with lung diseases like cystic fibrosis (7-10). Persistent infections lead to frequent antibiotic exposures and AMR emergence.

Ceftazidime/avibactam (CZA), ceftolozane/tazobactam (C/T), imipenem/cilastatin/relebactam (IMI/REL) are antimicrobials developed for treating multidrug-resistant (MDR) infections. These beta-lactam/beta-lactamase inhibitor combinations (BL/BLI) inhibit the transpeptidase activity of penicillin-binding proteins (PBPs) in the cell wall and block beta-lactam degrading enzymes (11-13). While studies have determined many possible mechanisms of resistance to these agents (14-20), multiple coexisting mutations in isolates muddle conclusions about which mutations are necessary for resistance. We present the evolution of coincident CZA, C/T, and IMI/REL resistance during CZA treatment of one infection and determine the resistance-driving bacterial mutations to inform treatment and drug development for resistant organisms.

Serial *P. aeruginosa* isolates cultured from a patient with a tracheostomy tube and a peritoneal infection were collected from the clinical microbiology lab of UPMC Children’s Hospital of Pittsburgh (IRB: STUDY21080186). Early isolates were from tracheal aspirate specimens and CZA treatment isolates were from a peritoneal infection. Ten isolates from eight sample dates were tested. The minimum inhibitory concentration (MIC) for CZA, C/T, and IMI/REL was determined using Liofilchem MIC Test Strips. DNA was extracted using DNAeasy Blood and Tissue Kit (Qiagen). Short read whole genome sequencing was performed to a minimum depth of coverage of 1.3 million reads by SeqCoast Genomics (Portsmouth NH), reads were trimmed with Trimmomatic and genomes were assembled using SPAdes (21, 22). Genome annotation was performed using Prokka and variant calling was completed using breseq v0.38 (23-26). A phylogenetic tree was created using RAxML v8.2.11 with the rapid bootstrap algorithm, 2421 random seed, and 1000 runs using core genome assemblies created with Panaroo (27-29) and visualized using iTOL (30).

The draft genome assembly of the ancestor (Isolate 1) was good quality consisting of 109 contigs with 6.867 Mbp found in contigs ≥ 10,000 bp and N50 of 172,009. All isolates were monophyletic with a maximum of 10 SNPs between them (Fig 1A). The only strains considered resistant by Clinical Laboratory Standards Institute criteria (31) were strains 7b and 8b with CZA and C/T MIC of > 256 μg/mL and 8 μg/mL for IMI/REL, (Fig 1B, 1C, and 1D). While “b” isolates differed from their date-paired “a” isolate by colony morphology and resistance pattern, phylogenetic analysis confirmed that they were descendants of the initial strain (Fig 1A).

**Figure 1.**
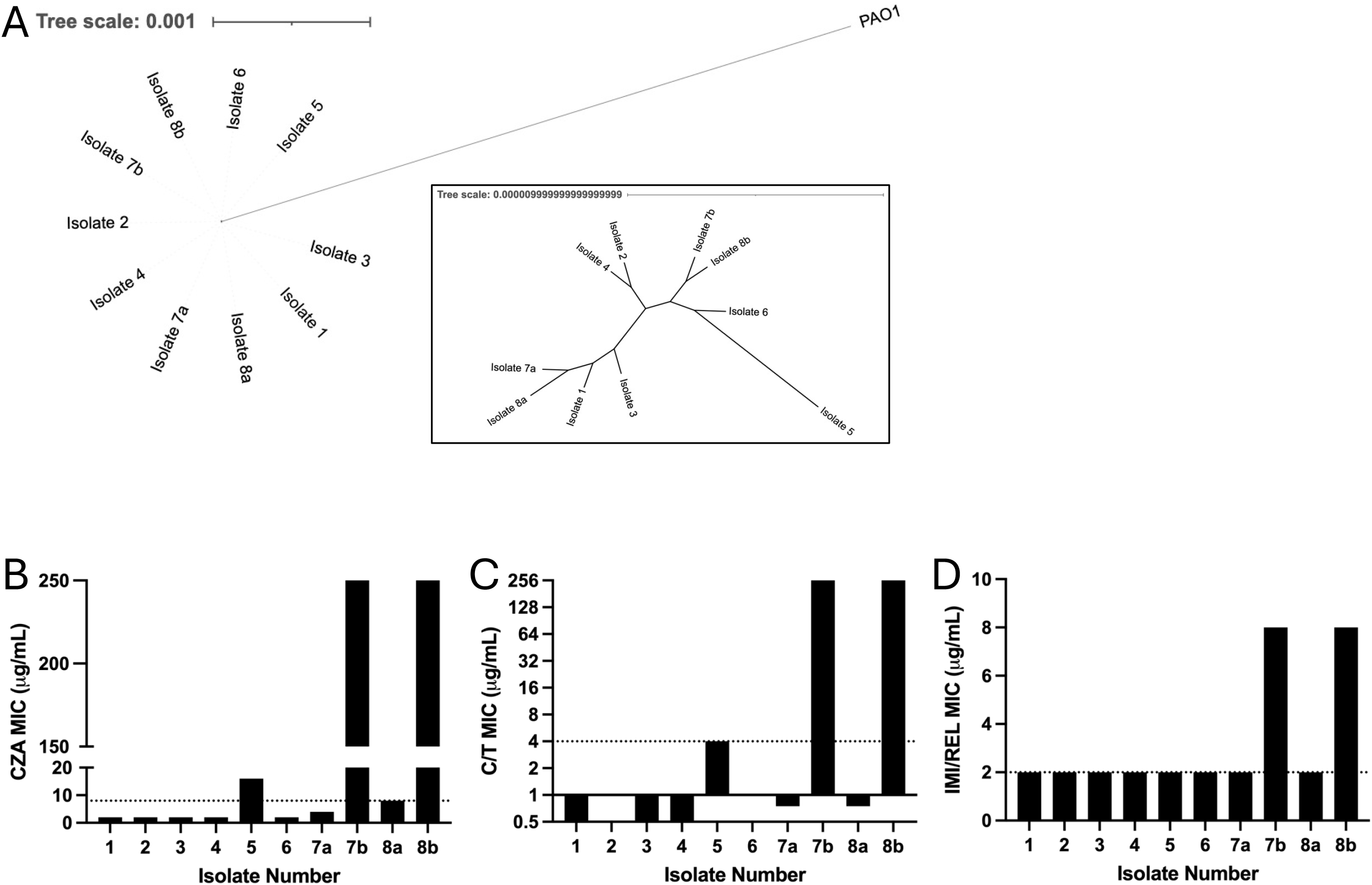
Isolates were all genetically linked and developed ceftazidime-avibactam and imipenem/relebactam resistance over time. (A) Phylogenetic tree demonstrating genetic relatedness of longitudinal isolates collected over time. Number indicates sample date and letter differentiate isolates from the same date. Inset shows zoom in of the isolates without PAO1 outgroup. (B-D) Minimum inhibitory concentrations (MIC) for isolates to (B) Ceftazidime/avibactam (CZA), (C) Ceftolozane/Tazobactam, and (D) Imipenem/Cilastatin/Relebactam (IMI/REL). Dashed lines indicate susceptibility breakpoints according to CLSI criteria.

To identify mutations that could explain the CZA, C/T, and IMI/REL resistance phenotype, we compared longitudinal isolates to the ancestor (23). We found three proteins with amino acid substitutions in CZA-, C/T-, and IMI/REL-resistant strains not present in susceptible strains: *ftsI* (R504C), encoding penicillin-binding protein 3 (PBP3), *bepR* (A95T), encoding an HTH-type transcriptional repressor (32), and *pvdS* (N73D), encoding a virulence regulator (33) (Fig 2A). As a transpeptidase for cell wall synthesis, the *ftsI* mutation relates directly to the mechanism of action of CZA, C/T, and IMI/REL. *bepR* is a homologue of *nalD*, a regulator of the MexAB-OprM drug resistance efflux pump. The mutation in *bepR* is outside the likely DNA-binding domain, residues 32-51, but could influence repressor function. We tested the ancestor and evolved isolates for efflux pump activity using an ethidium bromide (EtBr) assay (34). Efflux was increased in all isolates compared to ancestor, indicated by lower fluorescence. The resistant isolates, 7b and 8b, had equivalent efflux to 7a and 8a (Fig 2B), eliminating this as the primary resistance mechanism.

**Figure 2.**
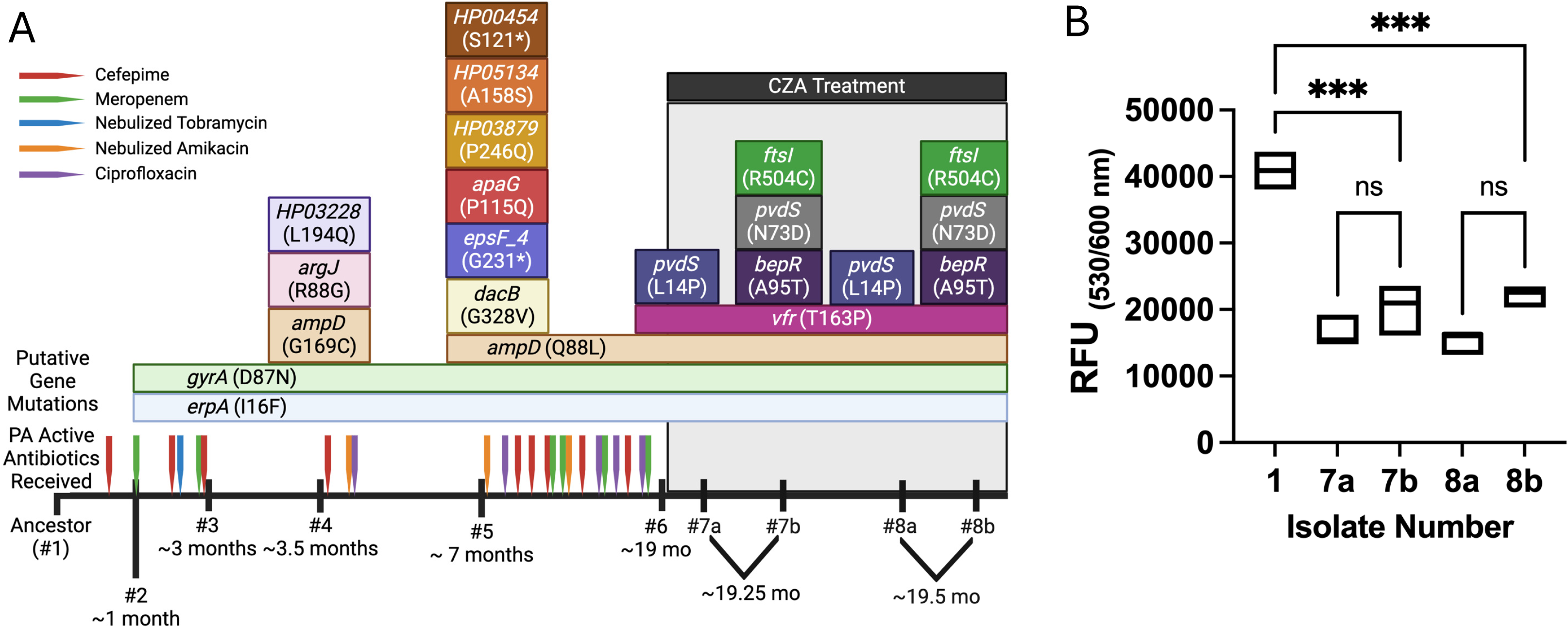
Putative mutations associated with CZA, C/T, and IMI/REL resistance do not confer increased efflux. (A) Timeline of accumulated mutations in clinical isolates over time. Number indicates sample date and letter differentiate isolates from the same date. Mutations are in colored boxes overlying the tick mark on the timeline for the isolate number. Gene names are indicated in italics with amino acid substitutions in parentheses. HP in mutation boxes indicates a hypothetical protein not annotated by Prokka. CZA treatment is shown as a black bar above the mutations. Times below the isolate numbers are relative to when the ancestor isolate (#1) was collected. Flags on timeline indicate treatment with *P. aeruginosa* active antibiotics prior to CZA treatment and color of flags indicate which antibiotic. (B) Ethidium bromide efflux assay was performed using all isolates normalized to bacterial density. Fluorescence readings were taken at one hour after addition of ethidium bromide and relative fluorescence units (RFU) is reported. Significance was tested using one-way ANOVA with Tukey’s multiple comparisons test. *** designates p < 0.001.

The development of coincident CZA, C/T, and IMI/REL resistance in this *P. aeruginosa* is best explained by the PBP3 mutation. Due to their mechanism of action, BL/BLIs provide selective pressure for mutations in penicillin-binding proteins and beta-lactamases. The PBP3 R504C substitution has previously been shown in beta-lactam-resistant isolates (17, 20, 35, 36) and respiratory isolates from persons with cystic fibrosis (37). This mutation was previously reported in IMI/REL-resistant strains, but other mutations were also present limiting causative conclusions (15, 17, 20). With minimal confounding mutations, we conclude that this PBP mutation is necessary cross resistance to novel BL/BLIs. While uncommon in *P. aeruginosa* compared to efflux and porin mutations (14, 38), PBP mutations represent a mechanism to eliminate the efficacy of multiple MDR-*P. aeruginosa* active antimicrobials.

The putative mechanism of cross resistance discovered here highlights the need to understand bacterial resistance mechanisms to novel antimicrobials to guide drug development and treatment for MDR organisms. A PBP mutation is necessary to confer CZA, C/T, and IMI/REL resistance, but may not be sufficient as there were other mutations identified in the resistant isolates. One confounder is that resistance developed in a genome in which AmpD was already mutated (Fig 2A). AmpD regulates AmpC beta-lactamase expression, and AmpD mutations have been associated with AmpC overexpression (39). It is possible that AmpC overexpression combined with the PBP3 mutation results in CZA, C/T, IMI/REL resistance, however this requires direct testing.

One limitation of this work is that the completeness of variant calling with short read assemblies can be limited by gaps in the assembled genome. However, the high quality of the genome assembly permitted identification of mutations that plausibly explain the cross resistance.

Our findings provide strong evidence that novel anti-pseudomonal antibiotics can drive cross resistance through PBP mutations. They highlight that changing antimicrobial class when resistance emerges to a novel BL/BLI during treatment may be necessary to avoid cross resistance. Additionally, they emphasize the value of longitudinal isolate whole-genome sequencing from infections to readily identify bacterial genetic causes of treatment failure. Lastly, they underscore that drug development should explore alternative antibiotic targets to prepare to circumvent future resistance to current antibiotics.

## Acknowledgements

The authors would like to acknowledge Nanami Kubota for creating the core genome file for phylogenetics using Panaroo. G.J.R is supported by the University of Pittsburgh School of Medicine Department of Pediatrics K12 (5K12 HD052892-17) and UPMC Children’s Hospital of Pittsburgh CHP Scholars Program. A.B.R is supported by NIH 1F31AI172279-01A1. D.V.T. is supported by funding from the Department of Medicine at the University of Pittsburgh School of Medicine. Additional funding provided by NIH 7U19AI158076-03 (V.S.C.). The funders had no role in study design, data collection and interpretation, or the decision to submit the work for publication.

